# Shifting balancing selection on a chromosomal inversion in island populations

**DOI:** 10.1101/2025.09.10.675386

**Authors:** Jun Ishigohoka, Miriam Liedvogel

## Abstract

Chromosomal inversions often underlie local adaptation and differentiation between populations. However, little is known about how selection acting on a balanced inversion shifts in different environments in the context of demographic events. Here, we combine population genomic approaches with extensive simulations to show that split events of island populations shifted a parameter of balancing selection on a polymorphic inversion. In Eurasian blackcaps (*Sylvia atricapilla*), an 8 Mb-long inversion has been maintained polymorphic in most populations across the specie’s distribution range by long-term balancing selection. The frequency of this inversion is consistently lower in island resident and continental resident populations compared to behaviourally ancestral continental migrant populations. Inference of population history shows that at least four groups of residents, specifically one continental population and three sets of populations on different island systems, originate from independent and simultaneous split events from one ancestral population. Such a demographic history indicates that the consistent reduction in the inversion frequency among these independent resident populations is unlikely due to shared stochastic events. Approximate Bayesian computation (ABC) applied to simulations of balancing selection under the blackcap demography indicates that the optimal frequency of the inversion in the regime of negative frequency-dependent selection, a type of balancing selection, became lower in island populations. These results highlight how parallel shifts in selection parameters in similar environments can contribute to genetic differentiation among populations at inversion loci.

## Introduction

Chromosomal inversions, a structural mutation with a reversed chromosomal segment, can play major roles in adaptation. Since the pioneering work by Dobzhansky (1947) in *Drosophila melanogaster*, inversion polymorphisms in multiple systems have been shown to be distributed along environmental gradient called “clines” (Ayala et al., 2013; Hager et al., 2022; Kapun et al., 2016; Lowry & Willis, 2010; Nosil et al., 2023). This spatial structure of inversion frequency has been considered to be the result of spatially varying selection (Ayala et al., 2013; Hager et al., 2022; Kapun et al., 2016; Kirkpatrick & Barton, 2006). Indeed, some traits under spatially varying selection have been shown to be associated with genotype of inversions with a clinal distribution (Hager et al., 2022; Huang et al., 2020; Kapun et al., 2016; Sanchez-Donoso et al., 2022; Todesco et al., 2020). Additionally, a number of characterised inversion loci are under some form of balancing selection and have been maintained polymorphic over a long period of time, and they are associated with large (often discrete) phenotypic variation within species (Chouteau et al., 2017; Hager et al., 2022; Jay et al., 2018; Knief et al., 2016; Küpper et al., 2015; Matschiner et al., 2022; Sanchez-Donoso et al., 2022; Tuttle et al., 2016).

Despite the widely recognised evolutionary importance of inversion polymorphisms, characterisation of the underlying mode of selection acting on inversions — spatially varying divergent selection with gene flow, overdominance (heterozygote advantage), and negative frequency-dependent selection — and measurement of parameter values in controlled conditions or natural populations are still limited (Ayala et al., 2013; Hager et al., 2022; Nosil et al., 2023). Furthermore, in empirical systems it is largely unknown how inversion clines are formed when populations expand, disperse, and/or split from their ancestral range to novel environments. One common challenge addressing these questions is the confounding by demographic history. Disentangling the selectively neutral effect of demography on a clinal pattern from selection on inversions requires modelling of the underlying demography finely tuned to the focal study system.

In this study, we investigate the evolutionary basis of geographically structured frequencies of a polymorphic inversion in the Eurasian blackcap *Sylvia atricapilla*. Blackcaps are common breeders across the species’ distribution range in Europe and Africa, and breeding populations (Fig. 1A) are characterised with differences in seasonal migration strategies (Berthold, 1991; Delmore et al., 2020; Shirihai et al., 2010). We previously identified an 8 Mb-long polymorphic inversion on blackcap chromosome 12 (inv_12_3) through genome scans for local genetic structure (Fig. 1A, (Ishigohoka et al., 2024)). Inv_12_3 contains mutations differentiated between two haplotypes (Fig. 1C), indicating that this polymorphism has been maintained for a long time by balancing selection (Ishigohoka et al., 2021). This focal inversion is present at different frequencies among populations with an apparent cline between the continental and island (as well as phenotypically migratory versus resident) populations (Fig. 1D) (Ishigohoka et al., 2024). We utilise this inversion system in blackcaps to address how selection acting on the inversion differs in independent populations after they split from the ancestral population.

**Figure 1:**
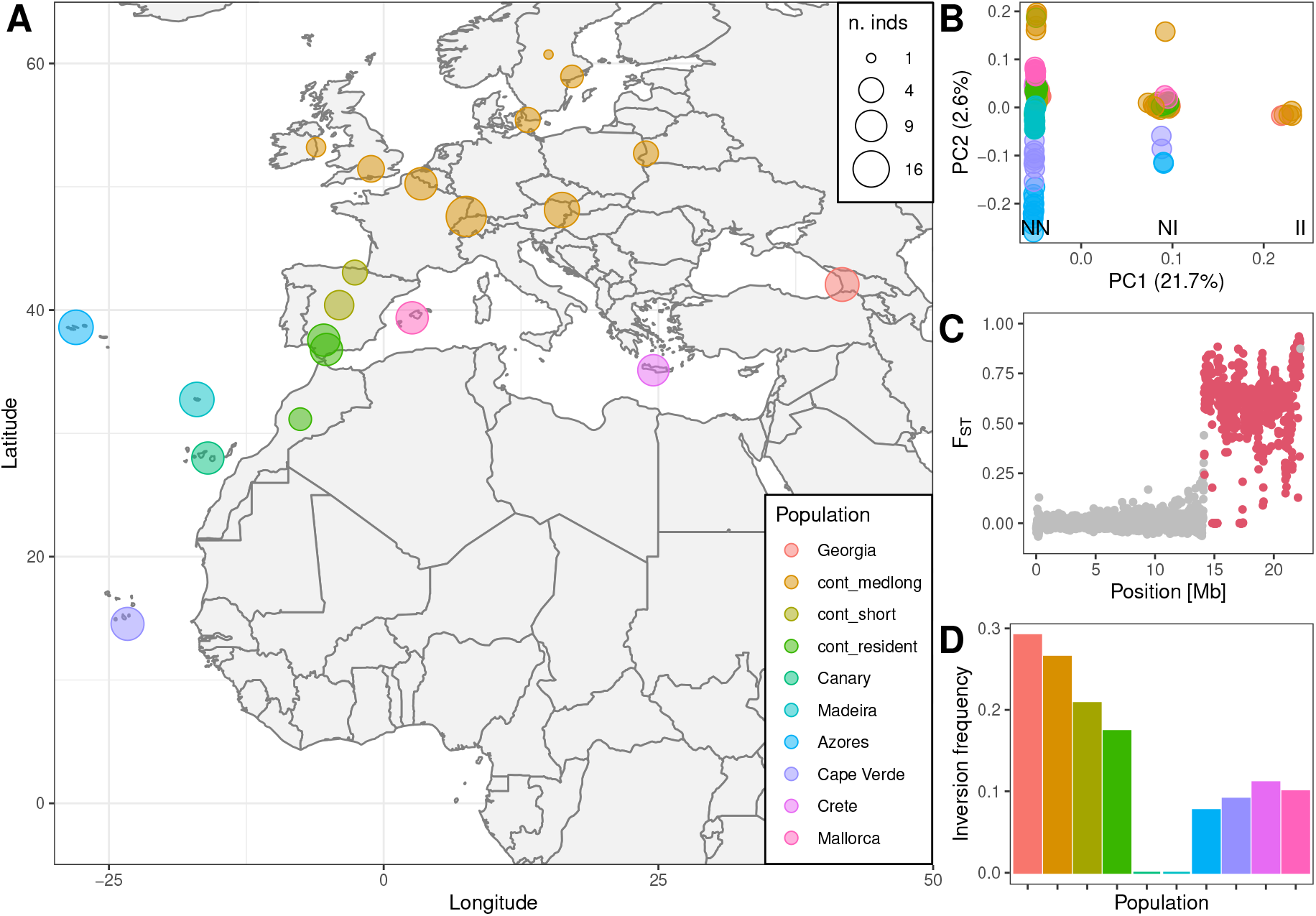
Inversion in blackcaps is under balancing selection. **A**. Geographic distribution of blackcap populations included in this study. **B**. Local population structure within the 8 Mb-long inversion of chromosome 12 based on PCA. The three clusters of individuals separated along PC1 correspond to their respective genotype at inv_12_3 (NN: normal/normal, NI: normal:inverted, II: inverted/inverted). **C**. F_ST_ in 10 kb windows between NN and II along chromosome 12. Red points highlight windows located within inv_12_3. **D**. The frequency of the inversion among the blackcap population.

## Results

The study entails two steps. First, we inferred the demographic history of the blackcap populations in focus here using whole-genome SNPs. This inference of neutral demography included identification of a demography model and estimation of demography parameters, which allowed us to simulate balancing selection under the inferred blackcap population history in the second step. Second, we characterised shifting balancing selection using simulation and observed inversion frequencies. We focused on two types of balancing selection: overdominance and negative frequency-dependent selection (NFDS). We excluded the possibility of divergent selection across the cline, because the frequency of inv_12_3 stays below 0.5 throughout our sampled populations (Fig. 1A, D) which covers most of the species breeding range (Shirihai et al., 2010). We identified a model from neutrality, overdominance, and NFDS, and estimated parameters.

### Demography inference reveal simultaneous and independent split of populations with lower inversion frequency

To characterise demographic history of blackcap populations, we used whole-genome resequencing data of 179 blackcaps covering their breeding range (Delmore et al., 2020; Ishigohoka et al., 2024). We first used three “model-free” methods for demography inference: `Stairway plot 2` (Liu & Fu, 2020), `MSMC2` (Malaspinas et al., 2016; Schiffels & Durbin, 2014; Schiffels & Wang, 2020), and `Relate` (Speidel et al., 2019). `Stairway plot 2` infers piecewise effective population size (N_e_) over time for a population based on the site-frequency spectrum (SFS) and is robust to high recombination rates (Ishigohoka & Liedvogel, 2025). `MSMC2` approximates the ancestral process along the genome in a structure called ancestral recombination graph (ARG) by sequentially Markovian coalescent (SMC) (Malaspinas et al., 2016; McVean & Cardin, 2005; Schiffels & Durbin, 2014; Schiffels & Wang, 2020). It estimates piecewise coalescence rate over time between multiple pairs of sequences from a population or from a pair of populations, which are used to obtain piece-wise N_e_ for each population and relative cross-coalescence rate (rCCR) between the population pair over time. `Relate` estimates the underlying ARG as a series of marginal trees (“genome-wide genealogies”) based on local distance matrices (Speidel et al., 2019), which are constructed by modelling observed haplotypes and ancestral haplotypes using a hidden Markov model (“chromosome painting” (Li & Stephens, 2003)). Piecewise coalescence rates over time epochs between all pairs of haplotypes are estimated from the inferred genealogies, which can be used to obtain N_e_ and rCCR over time as in `MSMC2`. `MSMC2` and `Relate` are sensitive to the presence of wide high-recombining regions in the genome, which is prominent in birds including blackcaps (Ishigohoka & Liedvogel, 2025). `Stairway plot 2` was run using folded SFS of each blackcap population. `MSMC2` was run using a low-recombining half of the genome of at most four individuals per population. `Relate` was run using a low-recombining half of the genome using all individuals. The inferences were qualitatively similar (Fig. 2A, B, Sup. Fig. 1) with consistency in one important aspect of the demography: all resident populations started to split simultaneously.

**Figure 2:**
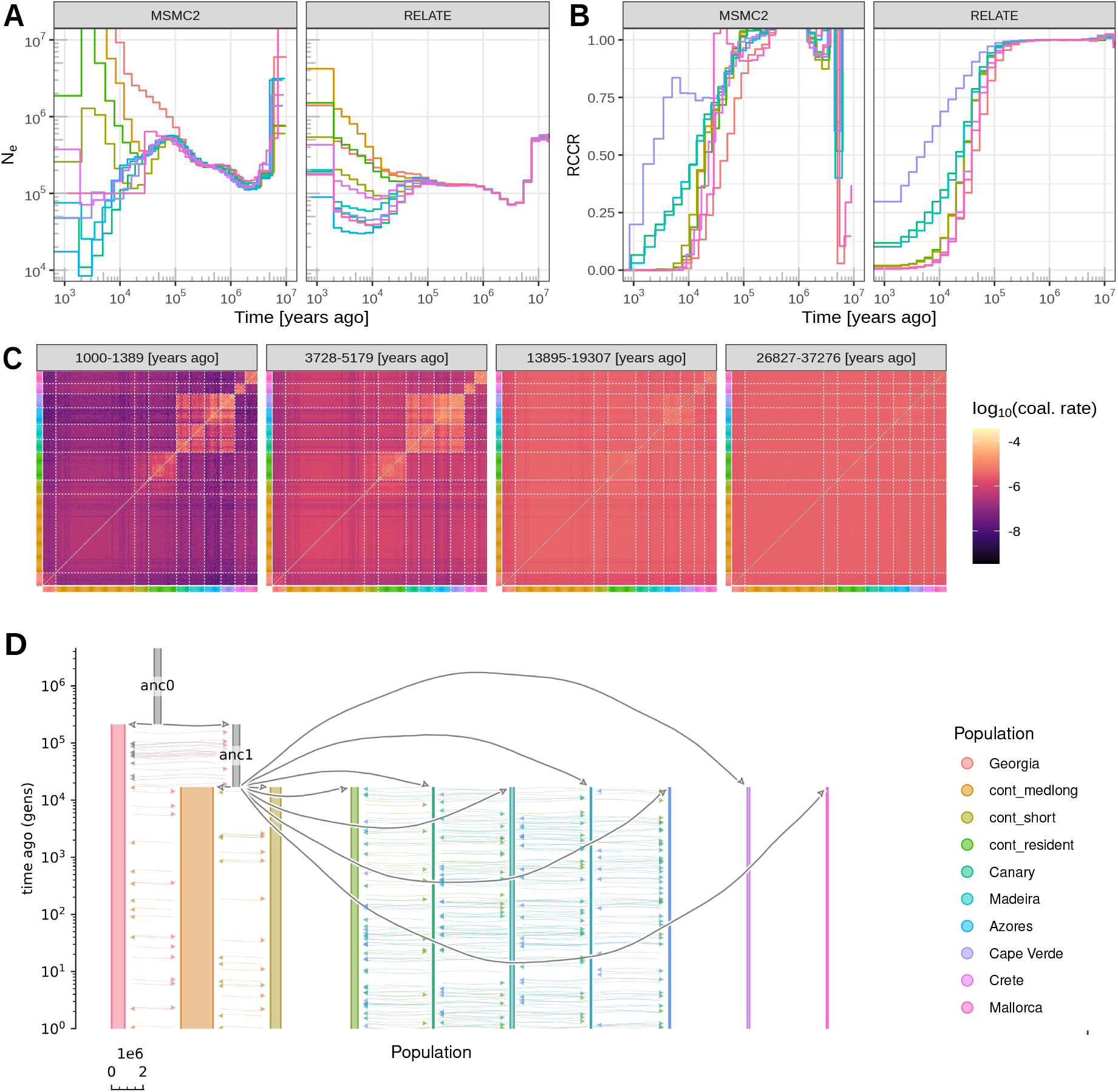
Demography inference. **A**. Historical effective population size (N_e_) of blackcap populations inferred with `MSMC2` and `Relate`. **B**. Relative cross-coalescence rate (RCCR) inferred with `MSMC2` and `Relate` for representative pairs of populations. Nine lines show RCCR between Azores and the other nine populations. **C**. Historical coalescence rate inferred with `Relate`. Each 358 x 358 heatmap shows coalescence rates between all 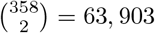 pairs of haplotypes from 179 individuals at the focal time epoch. The increase in coalescence rate in multiple “clusters” within the heatmap indicates that there were multiple independent isolation events across continental and island resident populations. **D**. Demography model inferred through approximate Bayesian computation (ABC) with `ABC-RF`. Parts of A and B are reproduced based on dataset from Ishigohoka & Liedvogel (2025).

To further investigate this aspect of population history (independent and simultaneous splits), we used genealogies inferred by `Relate` to summarise how coalescence rate changed over time (from present to past) for all pairs of haplotypes (Fig. 2C). We observed elevated coalescence rates in four distinct clusters of resident individuals: (1) continental resident population; (2) four Macaronesian island populations (Canary Islands, Madeira, Azores, Cape Verde); (3) Mallorca island resident population; and (4) Crete island resident population. These clusters do not merge with each other until the signals disappear (backwards in time). This pattern, in addition to N_e_ and rCCR, indicates that at least four independent split events occurred simultaneously to form both continental and island resident populations. Crucially, all populations in these four groups show reduced frequency of inv_12_3 (Fig. 1D). Such consistent shifts in frequency in the same direction are unlikely to be explained purely by neutral drift through demography, indicating that the parameters of balancing selection on inv_12_3 shifted in a consistent manner across all islands and southern end of the continental distribution range (inhabited by continental residents).

We also observed inconsistencies among demography inferences by the three methods in the following three aspects. First, the split of the population ancestral to the current Georgian population was inferred to be older than split events of other modern populations based on `Stairway plot 2` (Sup. Fig. 1) and `MSMC2` (Fig. 2A, B, left column), whereas `Relate` inferred that all modern populations including the Georgian population split around the same time (Fig. 2A, B, right column). Second, N_e_ of the continental resident population was consistent across the split event according to `Stairway plot 2` (Sup. Fig. 1), whereas `MSMC2` and `Relate` inferred an increase in N_e_ (Fig. 2A). Lastly, the estimated values of N_e_ and time differed among these three methods.

To solve these inconsistencies and estimate demographic parameters, we performed approximate Bayesian computation (ABC) using simulations for eight different models, consisting of combinations of three binary factors. The first factor was whether or not the split of the ancestral population of the current Georgian population was older than that to other modern populations. The second factor was whether or not the continental resident population increased in N_e_ after the split. The last factor was whether or not gene flow should be included in the model between some pairs of populations. We included symmetrical gene flow between Macaronesian island populations and between continental populations based on rCCR (Fig. 2B) and historical coalescence rate (Fig. 2C). By setting parameter ranges covering quantitative differences in N_e_ and time among inferences by `Stairway plot 2`, `MSMC2` and `Relate`, we covered the space of models and parameters as large as necessary and as small as possible. Each of these eight models were simulated using `ms` (Hudson, 2002) over 1 million times with randomly sampled parameters. We used `abcrf`, an implementation of ABC with random forest (Raynal et al., 2019), for model selection. The model with the older Georgian split, constant N_e_ for the continental resident population and non-zero gene flow was selected over other models with a posterior probability of 0.698 (Sup. Table 1). Parameter inference using `abcrf` revealed that the ancestral population of the current Georgian population split first ∼424,000 years ago (assuming a generation time of two years), and other populations split ∼34,000 years ago (Sup. Table 2).

### Negative frequency-dependent selection shifted in parallel across demographically-independent island populations

We hypothesised that parameters of balancing selection shifted in some populations (island populations or resident populations) by unknown common environmental factors. We tested this hypothesis by addressing two layers of questions. First, we asked what type of balancing selection is acting on inv_12_3 (we term them “models”). Specifically, we asked whether it is overdominance (heterozygote advantage, Fig. 3A) or negative frequency-dependent selection (NFDS, Fig. 3B), and we included a neutral scenario as a control. Secondly, we asked in what populations the parameters shifted (Fig. 3C, D); we term them “scenarios”. Specifically, we asked whether parameters shifted in resident populations (scenario 2), in island populations (scenario 3), or differently in all populations (scenario 4), in addition to a scenario where parameters do not change (scenario 1).

**Figure 3:**
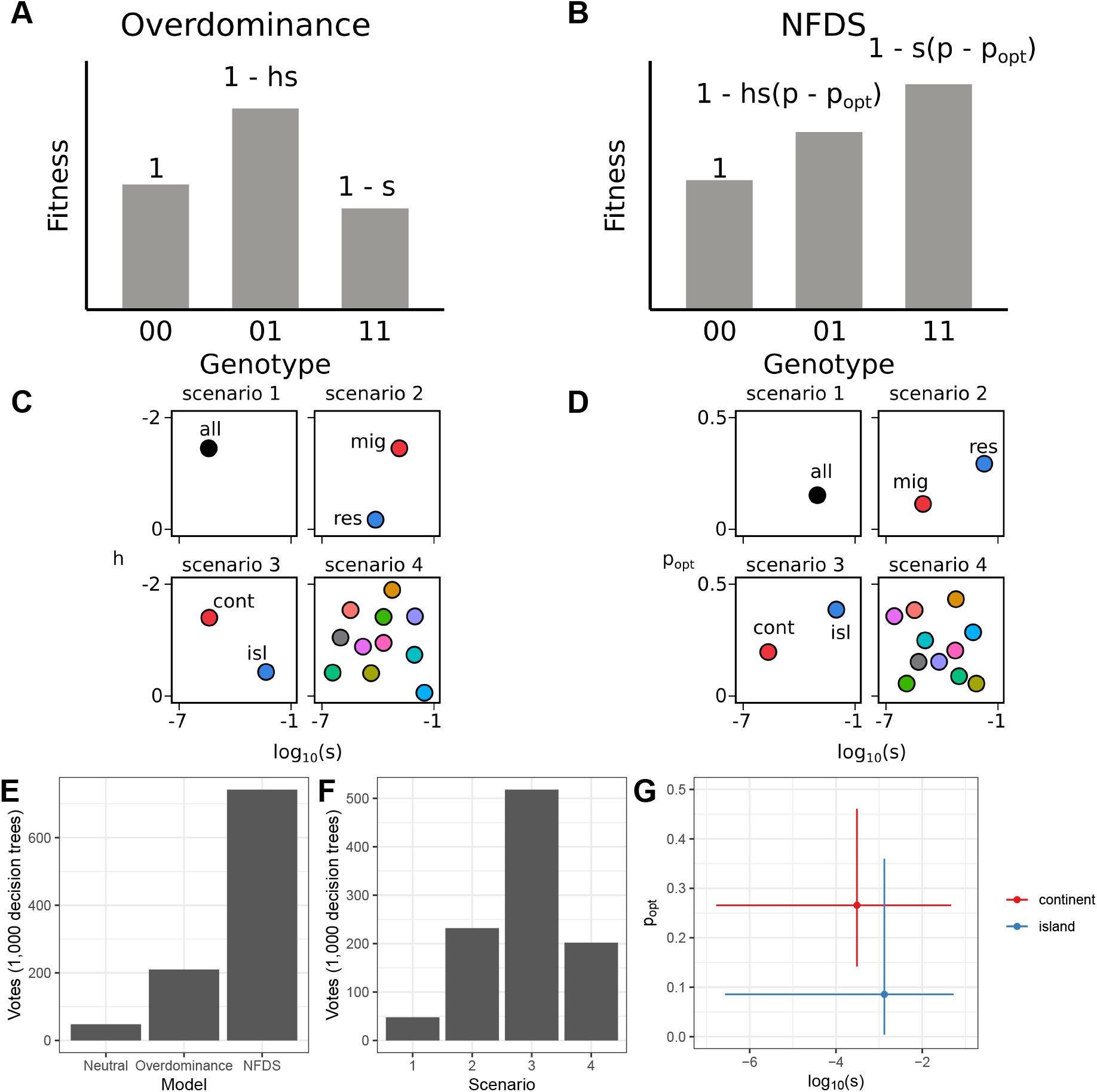
Shifting balancing selection. **A, B**. Two candidate models of balancing selection operating on the focal blackcap inversion inv_12_3. In the overdominance (heterozygote advantage) model (**A**), heterozygotes have higher fitness than homozygotes by the combination of two parameters, dominance (*h*) and selection (*s*) coefficients. In the negative frequency-dependent selection (NFDS) model (**B**), the fitness of the inversion depends on the difference between the inversion frequency *p* and the optimal frequency (*p*_*opt*_). We hypothesise that the lower frequency of the inversion on islands can be explained by different parameters of balancing selection (*h* and *s* in overdominance or *p*_*opt*_ and *s* in NFDS). **C, D**. Four scenarios of population difference in overdominance (**C**) or NFDS (**D**). In scenario 1, all populations share the same parameter value. In scenario 2, migratory (cont_medlong, cont_short, and ancestral population) and resident populations (cont_resident and all island populations) have different parameter values. In scenario 3, continental and island populations have different parameter values. In scenario 4, all populations have different parameter values. **E**. Model selection among neutral, overdominance, and NFDS with ABC-RF. **F**. Selection among four scenarios of NFDS with ABC-RF. **G** Parameter inference with ABC-RF. Points show the expectation, and error bars show 95% credibility interval.

To select models and scenarios, and to estimate parameters, we simulated selection on a locus for each scenario of each model over one million times in `SLiM` under the estimated blackcap demography, and performed ABC using `abcrf`. For each model, we sampled two parameters for simulation: selection coefficient (*s*) and dominance (*h*) for model 1 (overdominance); and selection coefficient (*s*) and optimal inversion frequency (*p*_*opt*_) for model 2 (NFDS). The two layers of questions (models and scenarios) were tested sequentially by two steps of model selection of ABC. For the first layer, the model for NFDS was selected over the other models with a posterior probability of 0.727 (Fig. 3E, Sup. Table 3). For the second layer, scenario 3 (parameters shifted in island populations) was selected over the other scenarios with a posterior probability of 0.619 (Fig. 3F, Sup. Table 4). Parameter inference revealed that the optimal frequency *p*_*opt*_ reduced in island populations to 0.086 from 0.266 in continental populations (Fig. 3G, Sup. Table 5). This result indicates that the frequency of inv_12_3 reduced in some populations because the optimal frequency of NFDS reduced in island populations.

## Discussion

Using approximate Bayesian computation (ABC), we revealed that the optimal frequency of negative frequency-dependent selection (NFDS) acting on inv_12_3, an inversion segregating across blackcap populations, shifted downwards in island populations. As a result of this shift, inv_12_3 is present at consistently lower frequencies in multiple independent island populations of blackcaps compared to continental populations. We simulated only scenarios with a discrete shift in the parameter of balancing selection. However, such a discrete shift may not be close enough to the mechanisms underlying the frequency of the inversion throughout the blackcaps’ distribution range. Although the frequency of inv_12_3 is higher in continental populations than in island populations, geographic heterigeneity of the inversion frequency exists also among the continental populations, reducing in frequency from northeast to southwest (Fig. 1D). In other words, the frequency of inv_12_3 follows a wide cline covering the entire breeding range of the species and also follows a cline with decreasing migratory distances exhibited by the population. The parameter values of balancing selection therefore may be shifting continuously along this axis. Although we aimed to capture this difference within continental populations in one scenario (scenario 2) considering different values of parameters in migrant and resident populations, an additional scenario (for both overdominance and NFDS) in which the focal parameter is expressed as a monotonic function of the location of populations along the cline axis should be carried out in a future study.

The target traits under balancing selection associated with the genotype of inv_12_3 were not identified in this study. One possibility is that inv_12_3 may harbour immune-related genes under balancing selection and they may have a different optimal frequency in island populations due to reduced parasite prevalence (Nieberding et al., 2006). Traits associated with inversions under selection may be identified after the finding of balanced polymorphism. For example, one inversion in the zebra finch was not associated with morphological traits or fitness components measured in the initial study (Knief et al., 2016), but was later identified to have an overdominance effect on sperm morphology and motility (Kim et al., 2017; Knief et al., 2017). Although quantifying morphological, behavioural, and fitness traits, including parasite prevalence, in wild-caught animals is not trivial, characterising the phenotypic and fitness effects of the inversion would be essential to understand the ultimate cause of the observed cline.

Inversions with clines in other systems are often associated with traits under spatially-varying selection (Ayala et al., 2013; Durmaz et al., 2018; Hager et al., 2022; Kapun et al., 2016; Kapun & Flatt, 2019; Kirkpatrick & Barton, 2006; Koch et al., 2021, 2022; Nosil et al., 2023). For example, an inversion in deer mice *Peromyscus maniculatus*, which is associated with forest and prairie ecotypes with different tail length (associated with climbing performance) and coat colour (crypticity in matching environment), is under divergent selection (Hager et al., 2022). Inversions in marine snails *Littorina saxatilis* are associated with weight, colour, and aperture size, which are diverged between ecotypes adapted to neighbouring intertidal habitats with opposing levels of pressure between crab predation and wave action (Koch et al., 2021, 2022). Inversions with latitudinal clines in *Drosophila melanogaster* are associated with multiple survival traits and are under selection by climatic factors (Durmaz et al., 2018; Kapun et al., 2016; Kapun & Flatt, 2019). An inversion in stick insects *Timema knulli* is under overall overdominance consisting of opposing effects of adaptation to a challenging host plant and survival (Nosil et al., 2023). In most of these inversions, the frequency difference between two sides of a narrow cline is so large that major allele switches across the cline. On the contrary to these opposing fitness effects across clines, inv_12_3 in blackcaps is maintained as a minor allele along the wide cline. This may represent a less extreme yet more general type of spatially varying balancing selection that does not involve flipping of the beneficial allele or genotype with the highest fitness.

## Materials and Methods

### Data and historical population structure

We used phased whole-genome resequencing (WGS) data of 179 blackcaps available from Ishigohoka et al. (2024). We revisited our demography inference with `MSMC2` using low-recombining half of blackcap genomes in Ishigohoka & Liedvogel (2025), as well as genome-wide genealogies of blackcaps inferred by `Relate` using low-recombining half of the blackcap genomes from the same study (Ishigohoka & Liedvogel, 2025). Historical population structure was estimated from genome-wide genealogies using the `RelateCoalescenceRate` module of `Relate`.

### Demography inference with ABC

To select a demography model and estimate demography parameters, we performed approximate Bayesian computation (ABC) with random forest using `abcrf` (Raynal et al., 2019), and `ms` (Hudson, 2002) as a simulator. We simulated eight scenarios consisting of the folloring three binary factors:

1. The population ancestral to the modern Georgian population split earlier than other modern populations (1) or it split with other populations simultaneously (2)
2. N_e_ of the continental resident population was constant from the ancestral population (1) or it increased (2)
3. There is no gene flow after population splits (1), or gene flow between some pairs of populations is included (2)

The prior distributions for parameters were: a log uniform between 1 *×* 10^5^ and 1 *×* 10^6^ for ancestral N_e_; a log uniform between 1 and 100 for the fold change in N_e_ in Georgian population, medium/long distance migrant population, and short distance migrant population (and continental resident population for models with an increase in N_e_ in the continental resident); a log uniform between 0.01 and 1 for fold change in N_e_ in island populations; a log uniform between 1 *×* 10^4^ and 1 *×* 10^5^ (generations) for split time of all populations (but the Georgians) for model with an earlier split of the Georgian population; a log uniform between the sampled split time of modern populations and 4 *×* 10^5^ (generations) for split time of the Georgian population in models with an earlier split of Georgian population. The prior distribution for gene flow (symmetric migration rate) was a log uniform between 1 *×* 10^−9^ and 1 *×* 10^−3^ (except for within Macaronesian islands), and seven (for models without an earlier split of Georgian population) or eight (for models with an earlier split of the Georgian population) values of gene flow were sampled: between the Georgian population and the population ancestral to all other populations; between Georgian and cont_medlong; between cont_medlong and cont_short; between cont_short and cont_resident; between Azores/Cape Verde and Canary/Madeira (“mig_mac”, mig for migratory and mac for Macaronesia); a log uniform from 1 *×* 10^−9^ to mig_mac between cont_resident and Macaronesian island populations; a log uniform from mig_mac to 1 *×* 10^−3^ between Azores and Cape Verde and between Canary and Madeira. Sampling of parameters and writing of `ms` commands with sampled parameters for different models were done in custom `AWK` scripts. In `ms` simulation, 1,000 independent loci with one mutation each were simulated, and the same numbers of individuals as in our empirical data set were sampled from these populations. The `ms` output was directly piped into an `AWK` script to compute 75 summary statistics: mean pairwise nucleotide difference *π* and Watterson’s *θ* for each population, and Weir and Cockerham’s estimator of F_ST_ for all pairs of populations. This simulation was performed 1,000,000 times for each of our eight models. For the empirical data, we randomly sampled 1,000 SNPs from autosomes and computed the summary statistics using the same script.

To perform model selection, a random forest was constructed using the `abcrf` function of the `abcrf` package. Model selection was performed using the `predict` method of the random forest. Cross validation was performed using the `err.abcrf` function of `abcrf` package. Goodness of fit was visualised using the `gfitpca` function of the `abc` package (Csilléry et al., 2012).

To estimate parameters, a regression random forest was constructed using the `regAbcrf` function of `abcrf` with a random forest of 1,000 trees. Parameters were estimated using the `predict` method of the regression random forest.

### Selection inference

To select a model of selection on inv_12_3, we performed ABC with random forest using `abcrf` using `SLiM` version 4.1 (Haller & Messer, 2022) as a simulator. We simulated one neutral model, one model for overdominance, and one model for NFDS. For the overdominance and NFDS models, we included four scenarios. In scenario 1, where all populations have the same parameters, a pair of parameters were sampled (*s* and *h* for overdominance and *s* and *p*_*opt*_ for NFDS). In scenario 2, where parameters differ between migrant and resident populations, two pairs of parameters were sampled and assigned to migrant and resident populations. In scenario 3, where parameters differ between continent and islands, two pairs of parameters were sampled and assigned to continental and island populations. In scenario 4, ten pairs of parameters were sampled, and they were assigned to different populations. For both overdominance and NFDS models, *s* was sampled from a log uniform distribution between 1 *×* 10^−7^ and 1 *×* 10^−1^. For the overdominance model, *h* was sampled from a uniform distribution between 0 and -2 (corresponding to expected inversion frequencies from 0 to 0.4). For the NFDS model, *p*_*opt*_ was sampled from a uniform distribution between 0 and 0.5. Sampling of parameters was done using custom `AWK` scripts.

We simulated a 1-bp locus representing inv_12_3 in `SLiM` version 4.0.1. We scaled the population size and time by a factor of 1,000. Assuming that the inversion was at the equilibrium frequency in the ancestral population, we introduced a marker mutation at the equilibrium frequency in the ancestral population (based on sampled *h* or *p*_*opt*_ for the ancestral population) in the initial generation. For the neutral model, we followed the same procedure as the overdominance model with *s* = 0. This leads to unrealistically high initial frequency for a neutral locus, hence the posterior probability for the neutral model might be overestimated. We implemented a rescaled version of the blackcap population history, and population-specific fitness effects were implemented within `mutationEffect` callback based on the genotype of the individual. At the final generation, we sampled the same numbers of individuals as our empirical data set, and recorded the inversion frequency. We ran 1,000,000 replicates of simulations per scenario.

We first selected a model (from neutral, overdominance, and NFDS), then selected a scenario (from scenarios 1-4) using the `abcrf` package. For both steps, we used the `abcrf` function to construct a random forest. A model and a scenario was selected using the `predict` method of the random forests. Cross validation was performed using the `err.abcrf` function of the `abcrf` package. Goodness of fit was visualised using the `gfitpca` function of the `abc` pakcage.

To estimate parameters, a regression random forest was constructed using the `regAbcrf` function of `abcrf` with a random forest of 1,000 trees. Parameters were estimated using the `predict` method of the regression random forest.

## Acknowledgments

This work was supported by the Max Planck Society (Max Planck Research Group grant MFFALIMN0001 to ML), and the DFG (project Nav05 within SFB 1372 – Magnetoreception and Navigation in Vertebrates (395940726) to ML). We thank Demetris Taliadoros for feedback on the study design and Alexander Jacobsen for helping with the implementation of simulation.

