## Supplementary Information for "Shifting balancing selection on a chromosomal inversion in island populations"

**Supplementary Table 1:** Model selection of demography models by ABC-RF.

| model | votes |
|---|---|
| 1_1_1 | 3 |
| 1_1_2 | 52 |
| 1_2_1 | 0 |
| 1_2_2 | 25 |
| 2_1_1 | 45 |
| 2_1_2 | 600 |
| 2_2_1 | 35 |
| 2_2_2 | 240 |

**Supplementary Table 2:** Parameter estimation of demography by ABC-RF. N_anc is $N_e$ of the ancestral population of all populations. N_N0_i ($i \in \{1, 2, 3, 5, 6, 7, 8, 9, 10\}$) is $N_e$ of population i relative to N_anc. T_geo is the splitting time of Georgian population from the ancestral population. T_split is the splittting time of all other populations. mig_i_j ($(i, j|i \neq j) \in \{1, 2, 3, 5, 6, 7, 8, 9, 10\}$)is the symmetrical migration rate (gene flow) between population i and j. 1 = Georgia. 2 = cont_medlong. 3 = cont_short. 5 = Canary. 6 = Madeira. 7 = Azores. 8 = Cape Verde. 9 = Mallorca. 10 = Crete.

| parameter | expectation | median | q2.5 | q97.5 |
|---|---|---|---|---|
| N_anc | 4.081e+05 | 4.253e+05 | 1.414e+05 | 9.495e+05 |
| N_N0_1 | 2.137e+00 | 1.981e+00 | 1.033e+00 | 6.521e+00 |
| N_N0_2 | 5.229e+00 | 4.799e+00 | 1.174e+00 | 3.750e+01 |
| N_N0_3 | 1.364e+01 | 1.530e+01 | 1.240e+00 | 9.178e+01 |
| N_N0_5 | 1.927e-01 | 1.964e-01 | 3.971e-02 | 8.624e-01 |
| N_N0_6 | 5.200e-01 | 5.824e-01 | 1.603e-01 | 9.733e-01 |
| N_N0_7 | 1.230e-01 | 1.193e-01 | 2.308e-02 | 7.256e-01 |
| N_N0_8 | 2.064e-01 | 2.068e-01 | 4.244e-02 | 8.786e-01 |
| N_N0_9 | 3.649e-01 | 3.553e-01 | 1.464e-01 | 8.920e-01 |
| N_N0_10 | 2.385e-01 | 2.350e-01 | 9.489e-02 | 6.078e-01 |
| T_geo | 1.891e+05 | 2.120e+05 | 4.145e+04 | 3.907e+05 |
| T_split | 1.854e+04 | 1.704e+04 | 1.018e+04 | 5.056e+04 |
| mig_0_1 | 1.174e-07 | 1.008e-07 | 1.239e-09 | 3.968e-05 |
| mig_1_0 | 1.174e-07 | 1.139e-07 | 1.207e-09 | 3.340e-05 |
| mig_1_2 | 1.823e-07 | 1.983e-07 | 1.270e-09 | 2.736e-05 |
| mig_2_1 | 1.358e-07 | 1.334e-07 | 1.224e-09 | 2.815e-05 |
| mig_2_3 | 5.528e-07 | 5.222e-07 | 1.304e-09 | 4.767e-04 |
| mig_3_2 | 4.610e-07 | 4.181e-07 | 1.380e-09 | 4.518e-04 |
| mig_3_4 | 2.046e-07 | 2.224e-07 | 1.275e-09 | 2.805e-05 |
| mig_4_3 | 1.910e-07 | 2.068e-07 | 1.202e-09 | 2.663e-05 |
| mig_4_5 | 4.489e-08 | 3.384e-08 | 1.159e-09 | 5.060e-06 |
| mig_4_6 | 4.345e-08 | 3.441e-08 | 1.113e-09 | 5.193e-06 |
| mig_4_7 | 4.629e-08 | 3.886e-08 | 1.098e-09 | 5.039e-06 |
| mig_4_8 | 5.322e-08 | 4.937e-08 | 1.138e-09 | 4.884e-06 |
| mig_5_4 | 5.647e-08 | 4.891e-08 | 1.151e-09 | 5.407e-06 |
| mig_5_6 | 2.984e-05 | 3.510e-05 | 9.701e-07 | 2.091e-04 |
| mig_5_7 | 3.252e-06 | 6.232e-06 | 3.634e-09 | 4.391e-05 |

| parameter | expectation | median | q2.5 | q97.5 |
|---|---|---|---|---|
| mig_5_8 | 3.692e-06 | 7.108e-06 | 4.741e-09 | 4.214e-05 |
| mig_6_4 | 5.918e-08 | 5.366e-08 | 1.157e-09 | 5.152e-06 |
| mig_6_5 | 3.014e-05 | 3.599e-05 | 1.868e-06 | 1.909e-04 |
| mig_6_7 | 3.557e-06 | 6.487e-06 | 4.653e-09 | 4.166e-05 |
| mig_6_8 | 3.702e-06 | 6.704e-06 | 4.478e-09 | 4.285e-05 |
| mig_7_4 | 4.997e-08 | 4.405e-08 | 1.127e-09 | 4.590e-06 |
| mig_7_5 | 3.482e-06 | 5.910e-06 | 8.183e-09 | 4.399e-05 |
| mig_7_6 | 3.561e-06 | 6.312e-06 | 5.950e-09 | 4.234e-05 |
| mig_7_8 | 3.547e-05 | 4.116e-05 | 2.353e-06 | 1.978e-04 |

**Supplementary Table 3:** Model selection of balancing selection models by ABC-RF.

| model | votes |
|---|---|
| neutral | 48 |
| OD | 210 |
| NFDS | 742 |

**Supplementary Table 4:** Model selection of shifting NFDS scenarios by ABC-RF.

| scenario | votes |
|---|---|
| scenario_1 | 48 |
| scenario_2 | 232 |
| scenario_3 | 518 |
| scenario_4 | 202 |

**Supplementary Table 5:** Parameter estimation of shifting NFDS by ABC-RF.

| parameter | expectation | median | q2.5 | q97.5 |
|---|---|---|---|---|
| s_cont | 3.061e-04 | 5.283e-04 | 1.670e-07 | 4.693e-02 |
| s_isl | 1.321e-03 | 3.009e-03 | 2.669e-07 | 5.369e-02 |
| p_opt_cont | 2.657e-01 | 2.520e-01 | 1.420e-01 | 4.610e-01 |
| p_opt_isl | 8.560e-02 | 6.800e-02 | 4.000e-03 | 3.600e-01 |

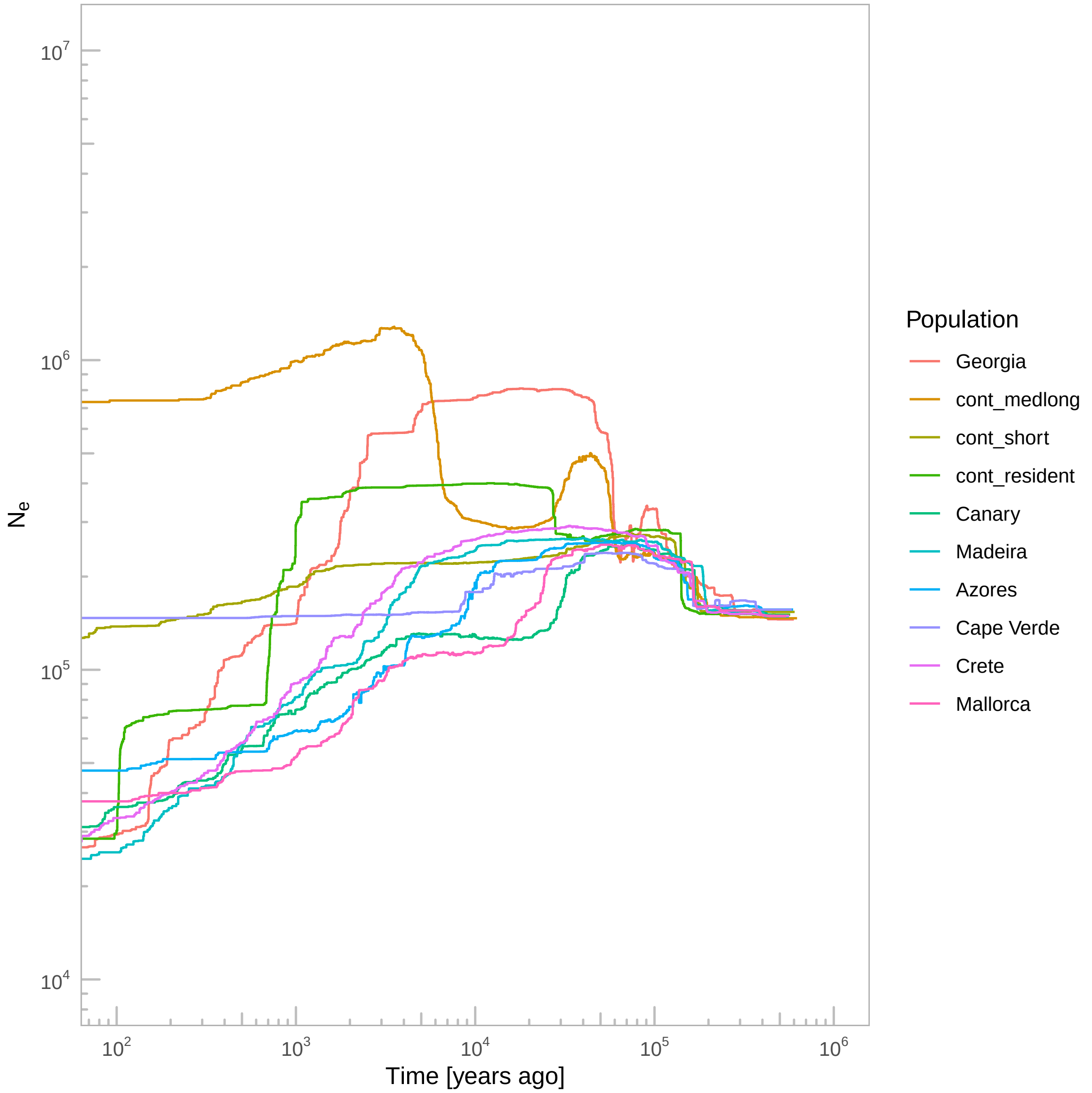


**Supplementary Figure 1:** Demography inference with `Stairway plot 2`.
